# rNMPID: a database for riboNucleoside Mono-Phosphates In DNA

**DOI:** 10.1101/2023.11.16.567417

**Authors:** Jingcheng Yang, Mo Sun, Zihan Ran, Taewhan Yang, Deepali L. Kundnani, Francesca Storici, Penghao Xu

**Affiliations:** State Key Laboratory of Genetic Engineering, School of Life Sciences, Human Phenome Institute, and Shanghai Cancer Center, Fudan University, Shanghai, China; Greater Bay Area Institute of Precision Medicine, Guangzhou, Guangdong, China; School of Biological Sciences, Georgia Institute of Technology, Atlanta, GA, USA; Department of Research, Shanghai University of Medicine & Health Sciences Affiliated Zhoupu Hospital, Shanghai, China; Inspection and Quarantine Department, The College of Medical Technology, Shanghai University of Medicine & Health Sciences, Shanghai, China

## Abstract

**Motivation:** Ribonucleoside monophosphates (rNMPs) are the most abundant non-standard nucleotides embedded in genomic DNA. If the presence of rNMP in DNA cannot be controlled, it can lead to genome instability. The actual positive functions of rNMPs in DNA remain mainly unknown. Considering the association between rNMPs embedment and various diseases and cancer, the phenomenon of rNMPs embedment in DNA has become a prominent area of research in recent years.

**Results:** We introduce the rNMPID database, which is the first database revealing rNMP-embedment characteristics, strand bias, and preferred incorporation patterns in the genomic DNA of samples from bacterial to human cells of different genetic backgrounds. The rNMPID database uses datasets generated by different rNMP-mapping techniques. It provides the researchers with a solid foundation to explore the features of rNMPs embedded in the genomic DNA of multiple sources, and their association with cellular functions, and, in future, disease. It also significantly benefits researchers in the fields of genetics and genomics who aim to integrate their studies with the rNMP-embedment data.

**Availability:** rNMPID is freely accessible on the web at https://www.rnmpid.org.

## 1 Introduction

Ribonucleoside triphosphates (rNTPs), the basic building blocks of RNA, are abundantly incorporated into DNA in the form of ribonucleoside mono-phosphates (rNMPs) by DNA polymerases due to their similarity to DNA nucleotides (Nick McElhinny *et al*., 2010; Brown and Suo, 2011). Previous studies suggested that the incorporated rNMPs constitute the most abundant non-standard nucleotides in the DNA of eukaryotic and prokaryotic genomes (Cerritelli and Crouch, 2016). Genome instability can result from failure to remove the genomic rNMPs, as the presence of rNMPs in DNA can alter the DNA structure, cause chromosomal fragility, and affect protein-DNA binding activity (Klein, 2017; Chiu *et al*., 2014; Williams *et al*., 2016). Furthermore, mutations in genes encoding any of the subunits of ribonuclease (RNase) H2, the main enzyme that initiates the rNMP removal, are found in the genotype of many patients affected by Aicardi-Goutières Syndrome (AGS) and Systemic Lupus Erythematosus (SLE), and in different types of cancer cells (Hiller *et al*., 2018; Aden *et al*., 2019; Moss *et al*., 2017; Williams *et al*., 2016). On the other hand, embedded rNMPs may also have physiological roles. For example, the abundant presence of rNMPs on the leading strand of DNA replication can guide mismatch repair in eukaryotic cells (Williams et al., 2013; Ghodgaonkar et al., 2013). Despite the threat posed to genomic integrity, abundant rNMP incorporation has persisted throughout the evolutionary scale, with millions of rNMPs in the human genome (Sassa *et al*., 2019). Therefore, there is still much to uncover about the functions and consequences of rNMPs embedded in DNA.

Over the last decade, researchers have devised multiple molecular biology techniques for mapping the genomic rNMPs, including ribose-seq (Koh *et al*., 2015), emRiboSeq (Ding *et al*., 2015), Alk-HydEn-seq (Clausen *et al*., 2015), RHII-HydEn-seq (Zhou *et al*., 2019), Pu-seq (Keszthelyi *et al*., 2015), GLOE-seq (Sriramachandran et al., 2020), and RiSQ-seq (Iida et al., 2021). Researchers have created over 200 libraries of embedded rNMPs in different species. These libraries help scientists understand how often rNMPs are included in DNA, the patterns of rNMP embedment, and how the rNMP presence relates to DNA metabolic functions in various organisms(El-Sayed *et al*., 2021; Balachander *et al*., 2020; Xu and Storici, 2021a; Kasiviswanathan and Copeland, 2011; Nick McElhinny *et al*., 2010; Xu *et al*.). Additionally, a series of bioinformatics tools have been developed to facilitate the mapping and analysis of rNMP incorporation in DNA (Gombolay and Storici, 2021; Xu and Storici, 2021b). With the help of sequencing-based rNMPs mapping technology, previous studies have utilized genome-wide rNMPs as indicators of replicative polymerases (Daigaku *et al*., 2015; Sriramachandran *et al*., 2020; Koyanagi *et al*., 2022). Other investigations have expanded our understanding of rNMP-incorporation functions by highlighting the direct interplay between ribonucleotide excision repair (RER) and topoisomerase 1 (Top1), two pathways for rNMPs removal, with the transcriptional processes in eukaryotic cells (Reijns *et al*., 2022; Hao *et al*., 2023). Exploring the role of rNMP embedment in relation to fundamental cellular processes such as replication and transcription, as well as its association with disease requires further study. This has made it a prominent area of research in recent years. To facilitate this goal for the scientific community, there will be an increased need to develop a comprehensive, multi-sourced database of rNMPs.

## 2 Methods

The rNMPID database is implemented by integrating more than ten published rNMP datasets derived from various species, including *Saccharomyces cerevisiae, Saccharomyces paradoxus, Schizosaccharomyces pombe, Chlamydomonas reinhardtii, Escherichia coli, Mus musculus*, and *Homo sapiens*. These rNMP datasets are constructed using four different rNMP-mapping techniques, ribose-seq (Koh *et al*., 2015), emRiboSeq (Ding *et al*., 2015), Alk-HydEn-seq (Clausen *et al*., 2015), and RHII-HydEn-seq (Zhou *et al*., 2019). Collected rNMP libraries are formatted in BEDGRAPH format and then converted to BigWig files for the genome browser. Afterward, by calculating the frequency and composition of rNMP embedment in various genetic elements, including genes, coding sequences, non-coding RNA, and other elements, we provide the researchers with valuable tools to study the association of rNMPs with such DNA elements and their role in various DNA metabolic processes. The count of rNMPs are normalized on the total number of rNMPs in the chosen rNMP library and the total nucleotide frequency from the background reference genome as previously described (Xu and Storici, 2021b). The rNMPID database is built using Rust (Matsakis et al., 2014), TypeScript (Bierman et al., 2014), PostgreSQL (Momjian et al., 2001) and additional libraries including Tokio (Tokio Team, 2023), SQLx (Launchbadge Team, 2023), Reactjs (Rawat and Mahajan, 2020), Plotly (Johnson et al., 2012), Ant Design (Ant Design Team, 2023), and JBrowse (Buels *et al*., 2016), etc. It comprises four different modules, namely Sample Analysis, Genome Browser, Download, and Resource (**Fig. 1A**). These modules offer the opportunity to conduct a comprehensive analysis and personalized visualization of rNMPs embedment in DNA.

**Fig. 1.**
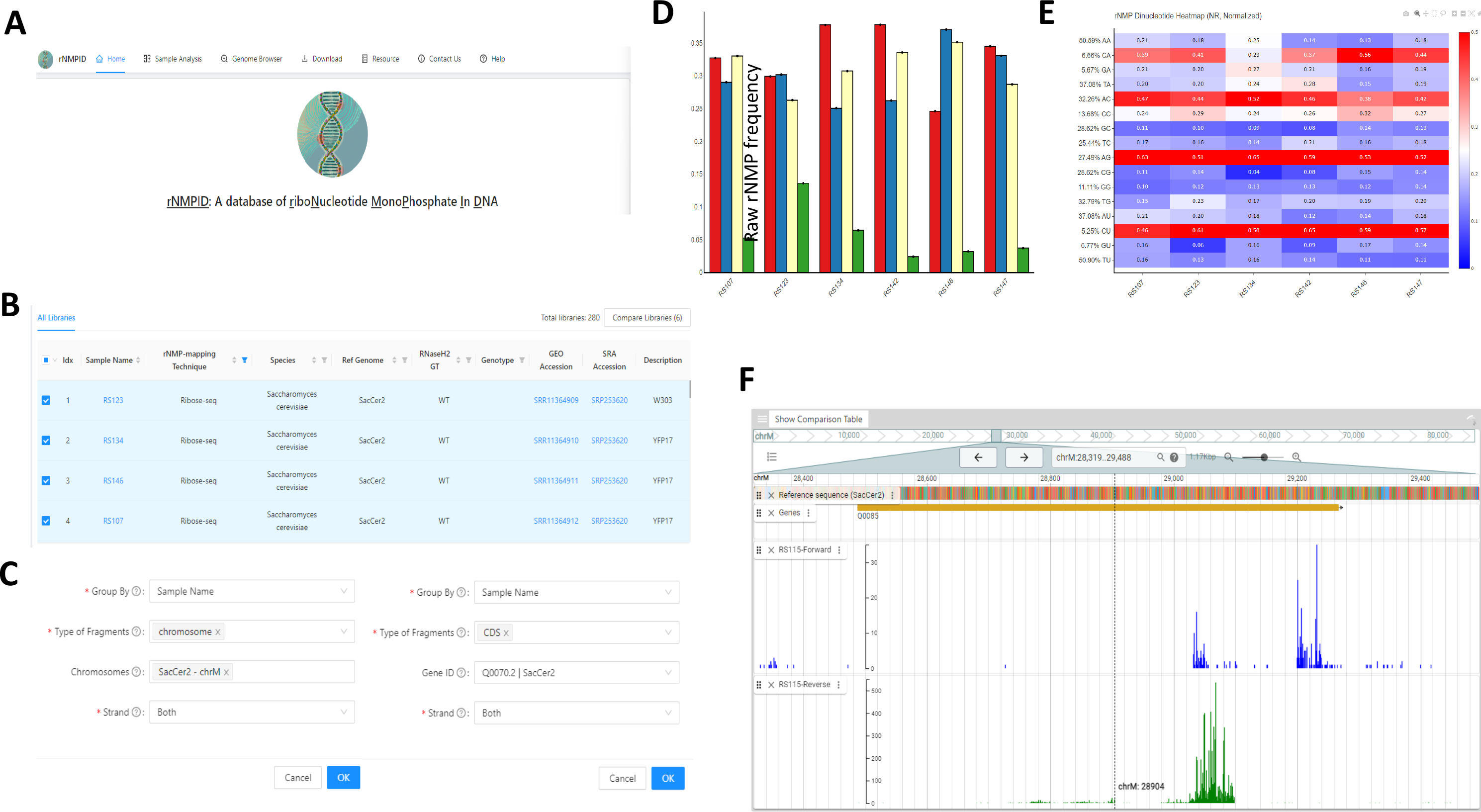
rNMPID database. (**A**) Homepage of the rNMPID database. (**B**) rNMP-sample metadata containing sample name, rNMP-mapping technique, species, reference genome, RNase H2 genotype, genotype, GEO accession, SRA accession, and description in Sample analysis module. (**C**) Users can select different types of DNA regions, and can also select the chromosome and the gene id in the Sample analysis module. (**D**) Example of bar plot showing the composition of the embedded rNMPs; red, rAMP, blue, rCMP, yellow, rGMP, green, rUMP in the selected libraries. (**E**) Example of heatmap showing the normalized frequency of dinucleotides composed of the incorporated rNMP (R: rA, rC, rG, or rU) and its upstream dNMP neighbor (N: dA, dC, dG, or dT) (NR). (**F**) An example of Genome Browser module with reference genome sequence, gene annotations, and rNMP-embedment sites.

## 3 Results

### 3.1 Sample Analysis module

The Sample Analysis module provides the users with handy tools to directly analyze the rNMP-embedment samples and reveal the rNMP-embedment characteristics. The metadata table in the module contains essential information about each collected sample, including sample name, rNMP-mapping technique, species, reference genome, RNase H2 genotype, genotype, GEO accession, SRA accession, and description (**Fig. 1B**). Users can select multiple rNMP samples of interests to perform analyses and comparisons. Moreover, the Sample Analysis module supports multiple query methods. Users can analyze the rNMP-embedment characteristics of the whole genome or focus on a single strand, on specific DNA fragments, genes (queried by Gene Name or Gene ID), and chromosomes to reveal the local rNMP-embedment patterns in the selected regions (**Fig. 1C**). By incorporating the RibosePreferenceAnalysis tool (Xu and Storici, 2021b), the Sample Analysis module performs various analysis on rNMP-embedment characteristics, including bar plots showing the raw and normalized frequency of the embedded rNMPs (**Fig. 1D**), heatmaps showing the rNMP-embedment compositions, and heatmaps showing the preferred dinucleotide patterns of rNMP embedment (**Fig. 1E**). By developing the Sample Analysis module, we devised to provide the researchers with handful tools to reveal the rNMP-embedment characteristics in the selected regions of interest of different samples, which can be easily integrated into their studies with the help of customizable visualizations.

### 3.2 Genome browser

To show the rNMP-embedment location and frequency on a single-nucleotide level, we incorporated the JBrowse Genome browser in our rNMPs database (Buels *et al*., 2016). The genome browser contains various tracks, including reference sequences, gene annotations, and rNMP location and frequency (**Fig. 1F**). Users can easily zoom in to their region of interest to see the frequency of rNMP embedment on each nucleotide and select multiple libraries to compare.

### 3.3 Resources

To assist users in performing their novel analysis on rNMP samples, we provided the free download of collected rNMPs sample data, reference genome, genome annotations, and formatted data showing rNMP-embedment characteristics used in the rNMPID database. We also gathered all the useful tools and studies in the Resources modules, including rNMP-mapping techniques, bioinformatics tools for rNMP-embedment analysis, and previous research related to the rNMP embedment in DNA.

## 4 Discussion and conclusion

The rNMPID database is a large-scale database. By initially integrating 280 rNMP-embedment libraries in six different species and four different rNMP-mapping techniques, rNMPID contains 258,516,657 unique rNMP-incorporation loci. This data amount is significantly larger than any other rNMP-related study.

The rNMPID database provides powerful data analysis and highly customizable visualization. We have incorporated state-of-the-art tools for the rNMP-embedment analysis in the rNMPID database. These tools offer researchers a comprehensive set of five distinct visualizations dedicated to examining rNMP composition and patterns in the Sample Analysis module. Each visualization is highly customizable, allowing users to modify parameters such as sample order, scale, and grouping methods to suit their specific research requirements. Additionally, the Genome Browser module and Download module empower users to perform in-depth investigations on the genomic region of interest.

Researchers can easily reveal the distinctive rNMP-incorporation characteristics across various species, cell types, and genotypes using rNMPID. We utilized rNMPID to replicate some key findings from a previous study of rNMP-incorporation characteristics within six wild-type and eight *rnh201*-null *S. cerevisiae* libraries (Balachander *et al*., 2020). In the Sample Analysis module, we initially selected these libraries from the metatable. Subsequently, we chose “RNH2 Genotype” and “nuclear DNA” as the criteria for “Group By” and “Type of Fragments” options. This analytical approach effectively reproduced our major findings. Notably, our results revealed that rC emerged as the most abundant rNMP within the nuclear DNA of both wild-type and *rnh201*-null cells, while rU appeared as the least abundant rNMP in *rnh201*-null cells (**Supplementary Fig. 1A, B**). Moreover, our examination of the genome browser revealed the presence of short nucleotide-repeated sequences, each displaying distinct patterns of rNMP enrichment. An illustrative example can be found at locus chrM: 63583 – 63651 within the RS156 library, where we selected RS156 in Genome Browser modules and chrM in the comparison table. Here, rNMPs were identified at the G-nucleotide position within the TAAGTA-repeated sequence on the forward strand and at the C-nucleotide position within the TACTTA-repeated sequence on the reverse strand (**Supplementary Fig. 1C**).

In summary, the rNMPID database is the first database of rNMP embedment in genomic DNA, which reveals rNMP-embedment characteristics, strand bias, and preferred rNMP patterns observed in the genome of different species in rNMP libraries generated using different rNMP mapping techniques, and in DNA samples of various genotypes. The rNMPID database provides the researchers with a solid foundation to explore the function of rNMPs embedded in genomic DNA and their association with DNA metabolic functions and potential disease.

## Acknowledgements

We thank Amazon Web Services (AWS) for funding and device support. We thank C. Gee from the Academic and Research IT for advice and technique support, and the Partnership for an Advanced Computing Environment (PACE) at the Georgia Institute of Technology for their research cyberinfrastructure resources and services. We acknowledge Y. Lee and S. Randhawa for critically reading the manuscript; and all members of the Storici Laboratory for assistance and feedback on this study.

