## Supplemental Figure1 for "rNMPID: a database for riboNucleoside Mono-Phosphates In DNA": Supplementary_fig1_final_bioaxiv.pptx

### Slide 1
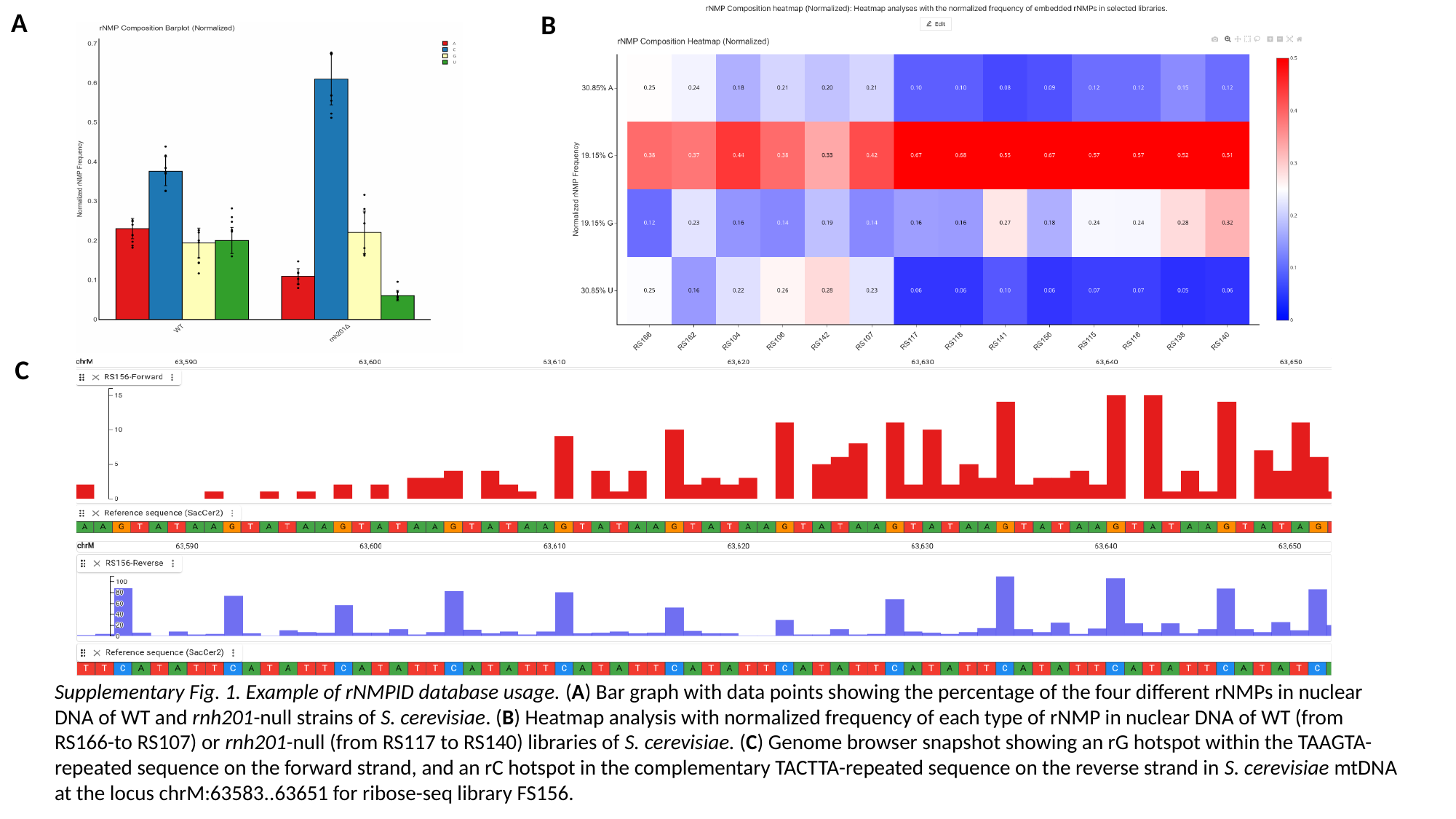

A
B
C
Supplementary Fig. 1. Example of rNMPID database usage. (A) Bar graph with data points showing the percentage of the four different rNMPs in nuclear DNA of WT and rnh201-null strains of S. cerevisiae. (B) Heatmap analysis with normalized frequency of each type of rNMP in nuclear DNA of WT (from RS166-to RS107) or rnh201-null (from RS117 to RS140) libraries of S. cerevisiae. (C) Genome browser snapshot showing an rG hotspot within the TAAGTA-repeated sequence on the forward strand, and an rC hotspot in the complementary TACTTA-repeated sequence on the reverse strand in S. cerevisiae mtDNA at the locus chrM:63583..63651 for ribose-seq library FS156.
